# Reconstitution of a flexible SYCP3-DNA fibre suggests a mechanism for SYCP3 coating of the meiotic chromosome axis

**DOI:** 10.1101/369439

**Authors:** Daniel Bollschweiler, Laura Radu, Jürgen M. Plitzko, Robert M. Henderson, Ioanna Mela, Luca Pellegrini

## Abstract

The synaptonemal complex (SC) keeps homologous chromosomes in close alignment during meiotic crossover. A hallmark of SC formation is the presence of its protein component SYCP3 on the chromosome axis. As SC assembly progresses, SYCP3 is deposited on both axes of the homologue pair, forming the lateral element (LE) in the tripartite structure of the mature SC. We have used cryo-electron tomography and atomic force microscopy to study the mechanism of assembly and DNA binding of the SYCP3 fibre. We find that the three-dimensional architecture of the fibre is built on a highly irregular arrangement of SYCP3 molecules displaying very limited local geometry. Interaction between SYCP3 molecules is driven by the intrinsically disordered tails of the protein, with no contact between the helical cores, resulting in a flexible fibre assembly. We demonstrate that the SYCP3 fibre can engage in extensive interactions with DNA, indicative of an efficient mechanism for incorporation of DNA within the fibre. Taken together, our findings suggest that, upon deposition on the chromosome axis, SYCP3 spreads by polymerising into a fibre that is fastened to the chromosome surface via DNA binding. The resulting layer of SYCP3 coating the chromosome axis might provide a structural basis for LE assembly in meiotic prophase.

---

At the heart of meiosis lies the exchange of genetic material between homologous parental chromosomes, which generates sexual diversity in the offspring. Correct and efficient recombination (crossover) between homologous chromosomes requires that they become physically aligned along their length. A supramolecular proteinaceous structure known as the synaptonemal complex (SC) is a critical participant in the process of crossover at meiotic prophase, by acting as a molecular zipper and linking the homologous chromosomes during recombination (1, 2). Despite considerable variations in the nature and sequence of its protein components, the SC structure shows a remarkable evolutionary conservation, consisting of lateral elements (LEs) that form on each paired chromosome and are linked by transverse filaments to a central element (CE) (3). Relatively little is known about the detailed structure of the SC, and how its dynamic architecture influences meiotic processes. Recent studies using super-resolution fluorescence have begun to provide important insights into the overall 3D structure of the SC and location of known factors (4-6).

SYCP3 is a major protein component of the LE (7). SYCP3 is recruited initially to the chromosome axis in leptotene, where it nucleates axial elements that spread and cover the entire length of the chromosome axis at pachytene, forming the lateral element of the tripartite structure SC (8). The molecular mechanism of LE assembly is unknown but is thought to result from the functional and physical interaction of SYCP3 with the SC component SYCP2 (9) and the meiotic hormads (HORMAD1 and 2 in human cells) (10). The three-dimensional architecture of the meiotic chromosome that is required for meiotic recombination results from the functional and physical interactions of SC components with meiotic cohesin complexes (11-16). A two-component system formed by a putative SYCP3 orthologue and a protein member of the Hormad family is present across organisms (for example, Red1 and Hop1 in yeast, ASY3 and ASY1 in plants) and might represent the structural core of the LE (17).

Correct SYCP3 function is essential for SC formation, chromosome synapsis and fertility (8, 18). *Sycp3* gene knockout causes infertility in male mice and reduced fertility in females, due to aneuploidy in oocytes and ensuing embryonic lethality (8, 19), and a mutation that renders SYCP3 defective causes human male infertility (20). Overexpression of SYCP3 has been reported in some types of cancer (21, 22). In SYCP3-null male mice, the LE and SC do not form and homologous chromosomes fail to achieve full synapsis (8, 16, 23). A two-fold increase in chromosome fibre length in SYCP3-deficient mice oocytes suggests a defect in chromosome organisation (19). Presumably, the LE must assemble on the existing structure of the chromosome axis, which is determined to a large extent by meiotic cohesins (13, 14, 24, 25).

SYCP3 folds into a highly elongated helical tetramer, where each chain forms antiparallel coiled-coil interactions and with two N- and C-tail protruding at each end of the helical core (26). A well-characterised property of SYCP3 is its ability to form filamentous fibres displaying transversal striations when overexpressed in mammalian cells (20, 27-29); this behaviour is mirrored by the ability of the recombinant protein to polymerise into similar, striated filamentous structures (26). Although the structural determinants driving SYCP3 polymerisation are presently unknown, self-assembly is critically dependent on sequence motifs in the N- and C-tails protruding from the helical core of the SYCP3 tetrameric structure (26, 28).

In addition to forming large filamentous structures, SYCP3 can interact with double-stranded DNA, via DNA-binding motifs located in its N-terminal tails (26). The presence of DNA-binding domains at either end of the elongated rod-like shape of the SYCP3 tetramer indicates that it can simultaneously interact with distinct segments of DNA (30). Single-molecule studies of the interaction of SYCP3 with DNA showed that DNA-bound SYCP3 molecules can form clusters that drive a limited degree of DNA compaction, in agreement with SYCP3’s structural role in determining LE architecture (30).

In this paper, we set out to elucidate how SYCP3 self-associates into filamentous fibres and how the SYCP3 fibres interact with DNA. Using a combination of cryo-electron tomography of the SYCP3 fibres and atomic force microscopy of SYCP3-DNA complexes we have obtained important new insights into SYCP3 function. We show that the regular higher-order structure of the SYCP3 fibre arises from a remarkably heterogeneous mode of association of individual SYCP3 particles, conferring plasticity to the fibre. Furthermore, we provide the first experimental evidence that polymeric SYCP3 fibres can engage in extensive interactions with DNA. Our results suggest that SYCP3 can coat the chromosome axis in a continuous structure containing both DNA-bound and DNA-free SYCP3 layers. We discuss the implications of this structural model for the function of SYCP3 and LE assembly in meiosis.

## RESULTS

### *In vitro* reconstitution of SYCP3 fibres

We had previously shown that recombinant human SYCP3 forms filamentous fibres that can be visualised by negative-stain Electron Microscopy (EM) on continuous carbon grids (26). The fibres show a periodic pattern of transversal, alternating light and dark bands, appearing at regular intervals of 22 nm. To determine whether SYCP3 fibre formation and its periodic appearance is an intrinsic property of SYCP3 in solution, we examined recombinant SYCP3 by cryo-electron microscopy. First, we established buffer conditions for controlled SYCP3 polymerisation, by determining that salt concentrations less than 300 mM KCl drive rapid SYCP3 self-association into a fibre (**Supplementary fig. 1**). We then exploited the salt-dependency of SYCP3’s aggregation state to prepare SYCP3 fibres in solution, which were preserved in vitreous ice.

Electron micrographs showed that SYCP3 fibres in vitreous ice maintained the same macroscopic organisation observed by negative-stain EM, consisting of transversal 22 nm-wide striations (**Fig. 1A** and **Supplementary fig. 2**). Different from the ribbon-like appearance of SYCP3 fibres prepared on continuous carbon grids, the SYCP3 fibre embedded in vitreous ice appeared as approximately cylindrical objects, with length in the micrometre scale and a diameter of the oval cross-section of 100 to 250 nm, indicating that the striation pattern extends three-dimensionally throughout the fibre (**Fig. 1A**). To confirm that fibre formation is a robust biochemical property of SYCP3 which persists under different preparation modes, we visualised the SYCP3 fibres by Atomic Force Microscopy (AFM). AFM analysis showed the presence of filamentous SYCP3 fibres that had a clear resemblance to those observed by EM, including the presence of the transverse 22 nm repeat (**Fig. 1B**).

**Figure 1.**
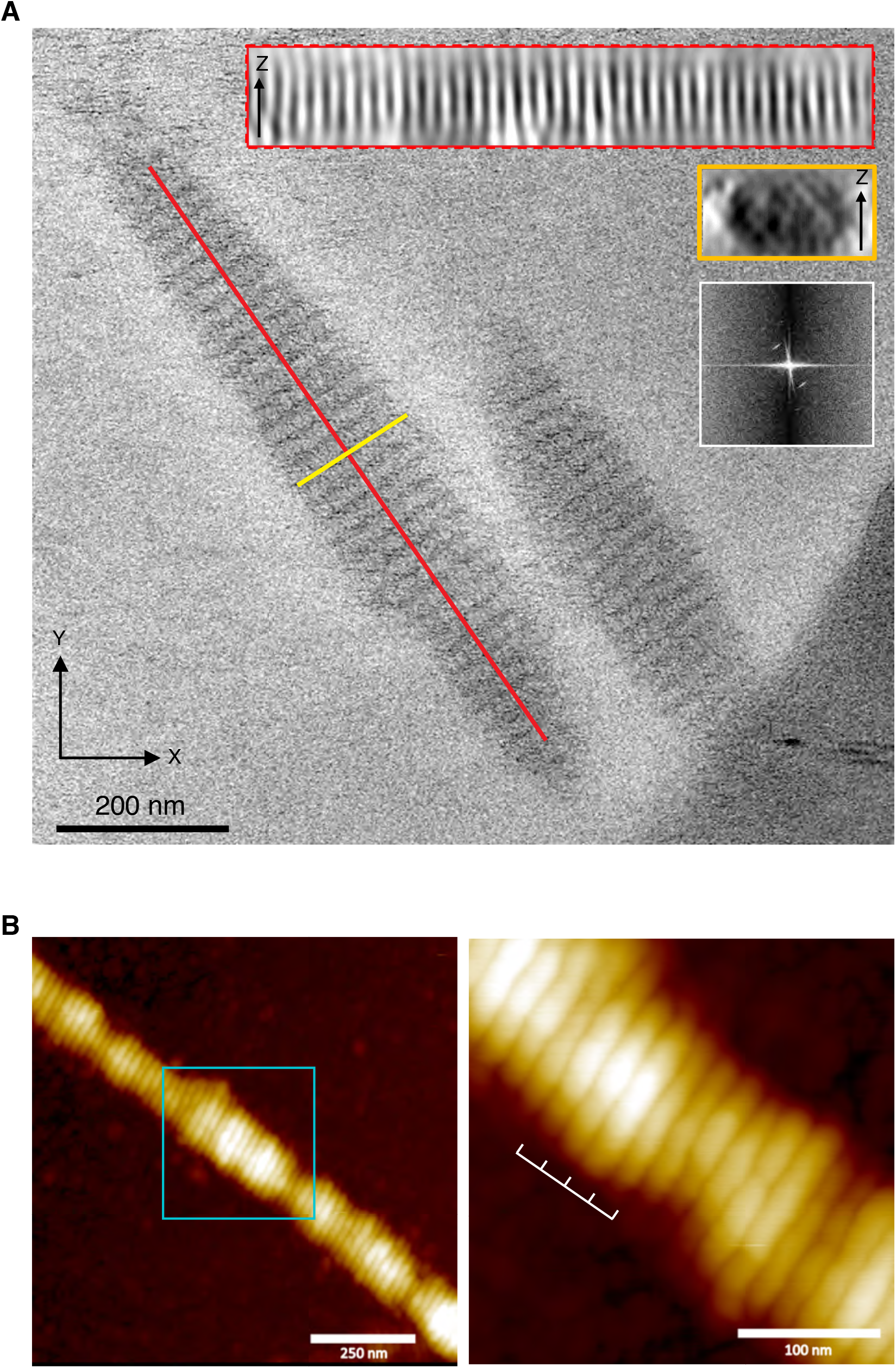
Structural analysis of the SYCP3 fibre. **A** Cryo-electron micrograph of the SYCP3 fibres, showing the regular pattern of transversal striations in the fibre. The cylindrically shaped fibres measure between 0.5 – 2 μm in length and 100 – 250 nm in diameter. The insets show two sections of the fibre, along and across its long axis (red and yellow boxes, respectively), and the Fourier analysis of the periodic striation of the fibre, revealing a layer line at a spatial frequency of 0.045 nm^-1^ (22 nm). **B** Atomic force microscopy shows that SYCP3 forms filamentous fibres that are similar to those observed by negative-stain electron microscopy (26) and by cryoEM (panel a). The right-hand panel shows a close-up view of the central portion of the fibre (cyan box), highlighting the presence of a transverse repeat of 22 nm.

As the cryoEM analysis had shown that the SYCP3 fibre maintains its characteristic periodic appearance in solution, we decided to investigate its three-dimensional architecture using cryo-electron tomography. A total of ten tomograms for native SYCP3 fibres were recorded under cryogenic conditions (**Supplementary movie 1)** and the tomographic volumes reconstructed (see Methods). The presence of a periodic striation pattern was confirmed by Fourier analysis, which revealed a corresponding layer line of spatial frequency of 0.045 nm^-1^ (22 nm).

### Sub-tomogram averaging of the SYCP3 fibre

To determine the structural basis for the homotypic SYCP3 interactions that underpin the three-dimensional architecture of the fibre, we performed sub-tomogram averaging. Due to the apparent dense packing of SYCP3 within the fibre and the low signal-to-noise typical of cryo-electron tomography, we did not attempt to pick individual SYCP3 particles for determination of initial subvolume coordinates. Instead, we chose the clearly identifiable 22 nm striation pattern as the guiding feature for subvolume coordinate selection. Thus, for each striation layer of the fibre, start and end points were selected manually, followed by automatic picking of intervening subvolume coordinate points (**Supplementary fig. 3**). Two sets of coordinate models were created this way. The first model, which was used to generate an initial reference structure *de novo*, had a large spacing of 44 nm (128 voxels) between each subvolume coordinate along the striation pattern. The second coordinate model featured a narrower spacing of 16.5 nm (48 voxels). This procedure was applied to the ten tomograms, resulting in 5,756 subvolume coordinates for set 1 and 46,051 for set 2.

A reference-free 2D classification of Z-projected subvolumes extracted from set 1 was then performed in Relion (31), to obtain 2D averages of the striation pattern and align all particles in the XY plane (**Fig. 2A**). The 2D-class averages indicated that each striation in the fibre contained a parallel array of elongated densities, which resembled the characteristic rod-like shape of the helical core of individual SYCP3 tetramers (26). The 2D averages further showed that the SYCP3 tetramers are aligned side-by-side in a direction parallel to the fibre long axis, and interact end-to-end to span the 22 nm repeat of the fibre.

**Figure 2.**
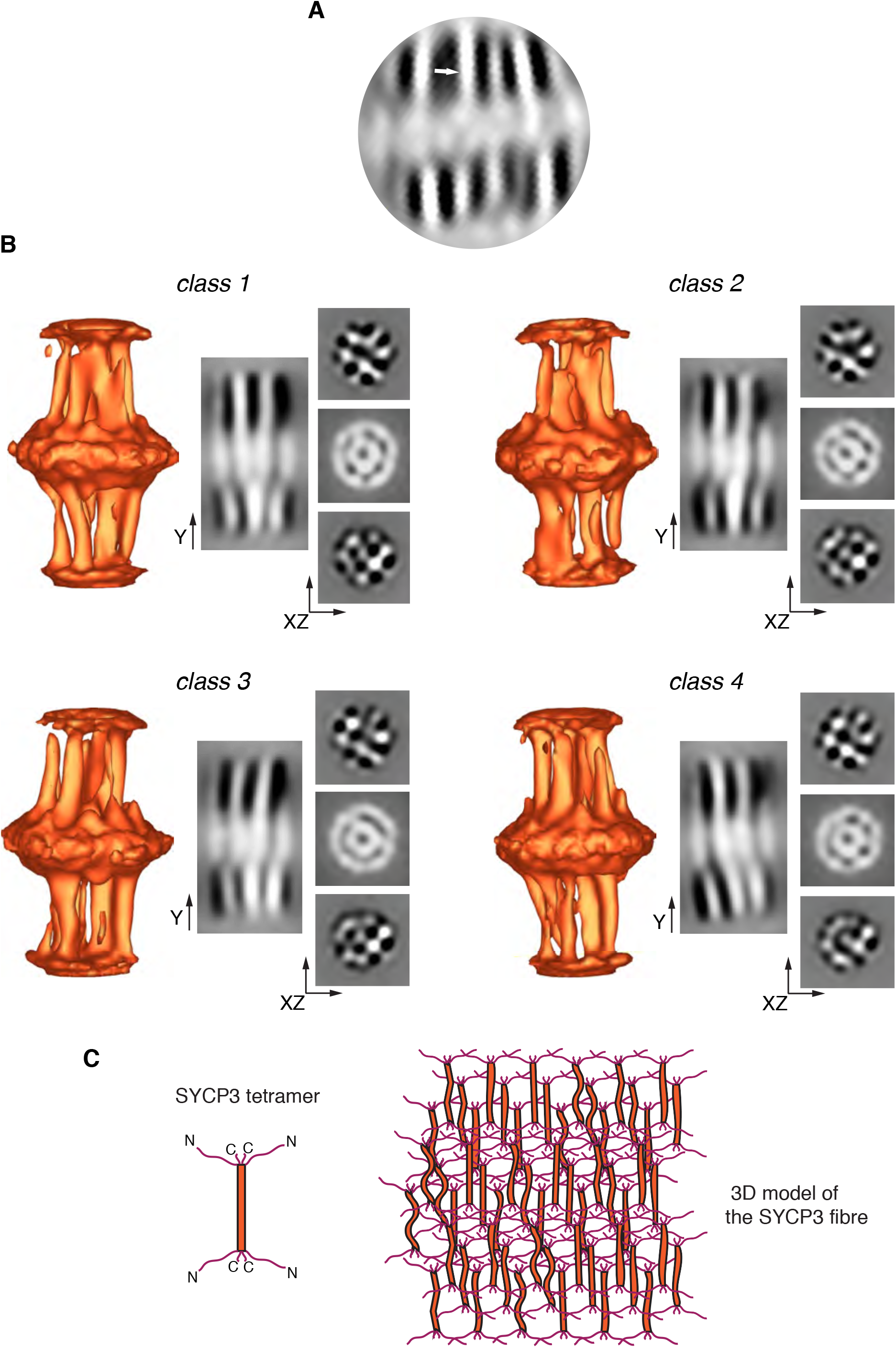
Final, refined class averages of the cylindrically masked SYCP3 fibre subtomograms. **A** 2D classification of projected subvolumes reveals clear evidence of density connecting adjacent striations (marked by an arrow), consistent with the shape of SYCP3’s tetrameric core structure (26). **B** Three-dimensional isosurface rendering of the four refined class averages. The classes had nearly equal distribution and tended to split subvolumes equally regardless of the number of specified classes (Supplementary figure 5). Isosurface sections along the Y axis and in the XZ plane are shown next to each isosurface rendering. The Y-axis sections through a three-particle region of the class averages highlight variations in conformation and spacing between individual SYCP3 particles. XZ cross-sections through the inter-striation region of all class averages show an approximately rectangular pattern of SYCP3 particles, but with a high degree of heterogeneity in particle arrangement between classes. **C** Schematic drawing for the three-dimensional arrangement of SYCP3 molecules in a fibre, based on the result of the cryo-electron tomography. The association between adjacent SYCP3 molecules are mediated exclusively by their intrinsically disordered tails, yielding a highly flexible polymeric assembly.

To gain further insight into the three-dimensional organisation of the SYCP3 fibre, we performed a maximum-likelihood 3D classification using Relion (31). Exploiting the parallel orientation of the fibres to the grid surface, the subvolumes were aligned based on their projections in the X,Y plane of the fibre. The alignments were then used as the starting point for a rotational 3D alignment. This procedure resulted in a low-resolution *de novo* structure which served as the initial reference model for masked 3D classification and 3D auto-refinement using the second, larger dataset (**Supplementary fig. 4**). The final class averages derived from the 3D classification revealed a remarkable degree of structural heterogeneity in the arrangement of SYCP3 molecules within the fibre **(Fig. 2B).** Importantly, further hierarchical 3D classification did not yield distinct, better-resolved models of the averaged sub-volumes (**Supplementary fig. 5**) indicating that a lack of long-range 3D order is an intrinsic architectural feature of the SYCP3 fibre.

Sub-tomogram averaging showed that individual rod-shaped SYCP3 tetramers are arranged in a parallel fashion along the long axis of the fibre. Occasionally, a kink or a bend is detected within a particle, in agreement with our previous observation of conformational flexibility in the middle of SYCP3’s helical core (26). Despite lack of side-by-side contacts between adjacent helical cores, the SYCP3 particles remain in vertical register within a layer. Thus, all contacts between SYCP3 particles appear to be mediated by the N- and C-extensions, taking place within the dense striations transversal to the fibre (**Fig. 2C**). A striking result of the 3D classification is the lack of geometric regularity in the planar arrangement of neighbouring SYCP3 particles, as highlighted by the XZ section of the 3D classes; the sections show only a loosely rectangular arrangement of SYCP3 particles that is rapidly lost beyond 2-3 repeating units. The implications of this finding for SYCP3 function are analysed in the Discussion.

### AFM analysis of SYCP3-DNA fibres

Alongside its ability to self-associate into a filamentous fibre, SYCP3 possesses a DNA-binding activity, mediated by the amino acid sequence N-terminal to its helical core. The combination of these two biochemical properties is likely to form the basis for SYCP3’s role in lateral element formation. Single-molecule experiments had provided evidence of DNA-dependent clustering by a C-terminally truncated version of SYCP3 that is defective in fibre formation (30). However, direct evidence that the SYCP3 fibre is competent to interact with DNA has been lacking.

To investigate its potential mode of DNA binding, we prepared samples of SYCP3 for AFM in the presence of plasmid DNA. The AFM images provided striking evidence of DNA incorporation into the SYCP3 fibre (**Fig. 3 and Supplementary fig. 6**). The regular and continuous presence of DNA loops projecting from the protein core along the fibre length indicates that an established mechanism of protein-DNA interaction operates within the proteinaceous core of the fibre. The ability of full-length SYCP3 to polymerise in a fibre depends critically on N- and C-terminal sequences flanking its helical core (26), and homotypic interactions were also found to be important for SYCP3-DNA fibre formation: SYCP3 1-230, lacking six residues at its C-terminus that are required for self-association but retaining DNA-binding ability, did not form extended fibres in the presence of DNA (**Fig. 4A**). Instead, clusters of DNA-bound SYCP3 1-230 were detected, in agreement with its behaviour in single-molecule optical tweezer experiments (30). DNA-bound fibres were also observed in the presence of plasmid DNA that had been linearised, but were of more limited extent (**Fig. 4B, C** and **Supplementary fig. 7**). The shorter length of the fibre when bound to linear DNA suggests that fibre growth is assisted by a degree of DNA supercoiling, probably because it facilitates DNA bridging by SYCP3.

**Figure 3.**
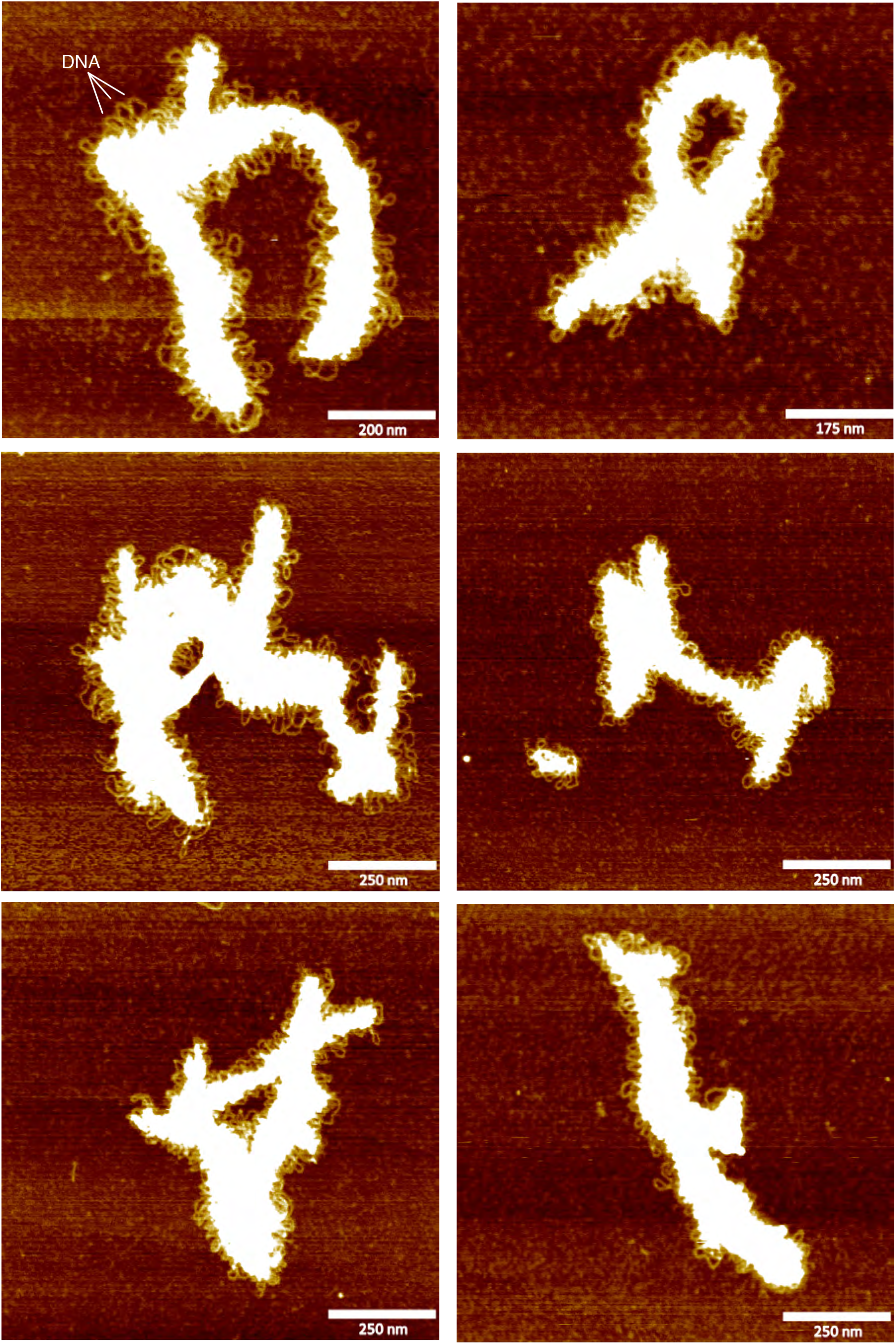
Atomic force microscopy of the DNA-bound SYCP3 fibre. Representative images of the fibrous structures formed by full-length SYCP3 in the presence of circular plasmid DNA. The DNA is visible as loops protruding from the protein core of the fibre.

**Figure 4.**
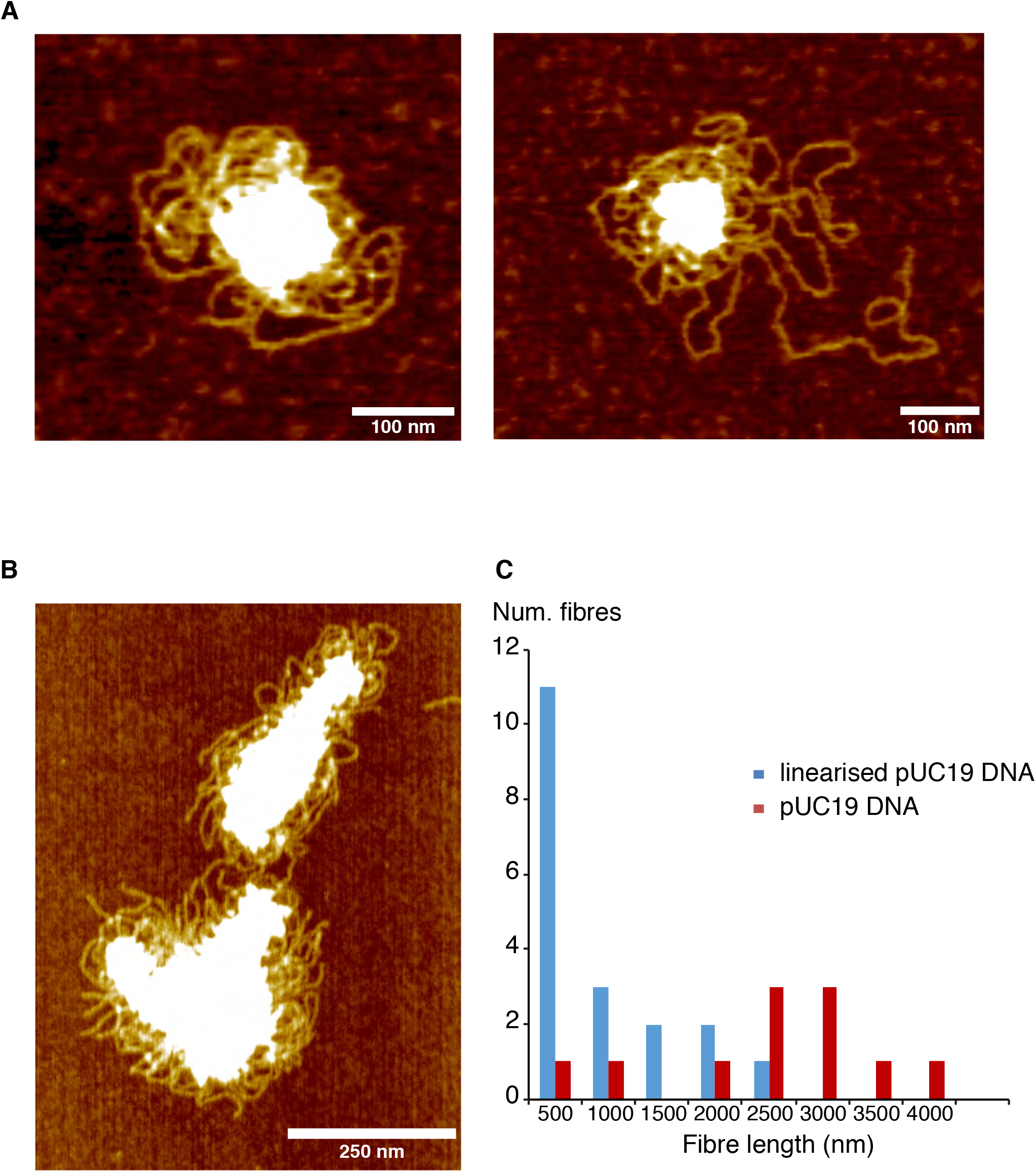
Observed changes in fibre appearance caused by modifications to the protein or the DNA. **A** AFM of C-terminally truncated SYCP3 1-230 in the presence of circular plasmid DNA (pUC19). The truncated protein tends to form clusters on the DNA, and no extended DNA-bound fibres are observed. **B** AFM of SYCP3 in the presence of linearized plasmid DNA. The left-side panel shows a representative image of SYCP3 bound to linearised plasmid DNA. On the right, the histogram distribution of fibre length in samples of SYCP3 formed in the presence of circular or linearized plasmid DNA.

Inspection of the DNA-bound SYCP3 fibres showed that fibre height was not uniform, but showed variation along the fibre length, ranging from 5 to 30 nm (**Fig. 5A, B**). A similar height distribution was observed for DNA-free SYCP3, with peaks at 10 and 20 nm (**Fig. 5B**) suggesting the presence of SYCP3 fibres consisting of two layers in thickness. Conversely, the height distribution of the fibre for the SYCP3 1-230 protein was sharper and clustered at around 10 nm (**Fig. 5B**). The greater height of the fibres formed by the full-length protein, in the presence or absence of DNA, might reflect its ability to form multi-layer assemblies, whereas the 10 nm layer of the SYCP3 1-230 protein might represent the height of the DNA-bound layer, due to the inability of the truncated protein to self-associate.

**Figure 5.**
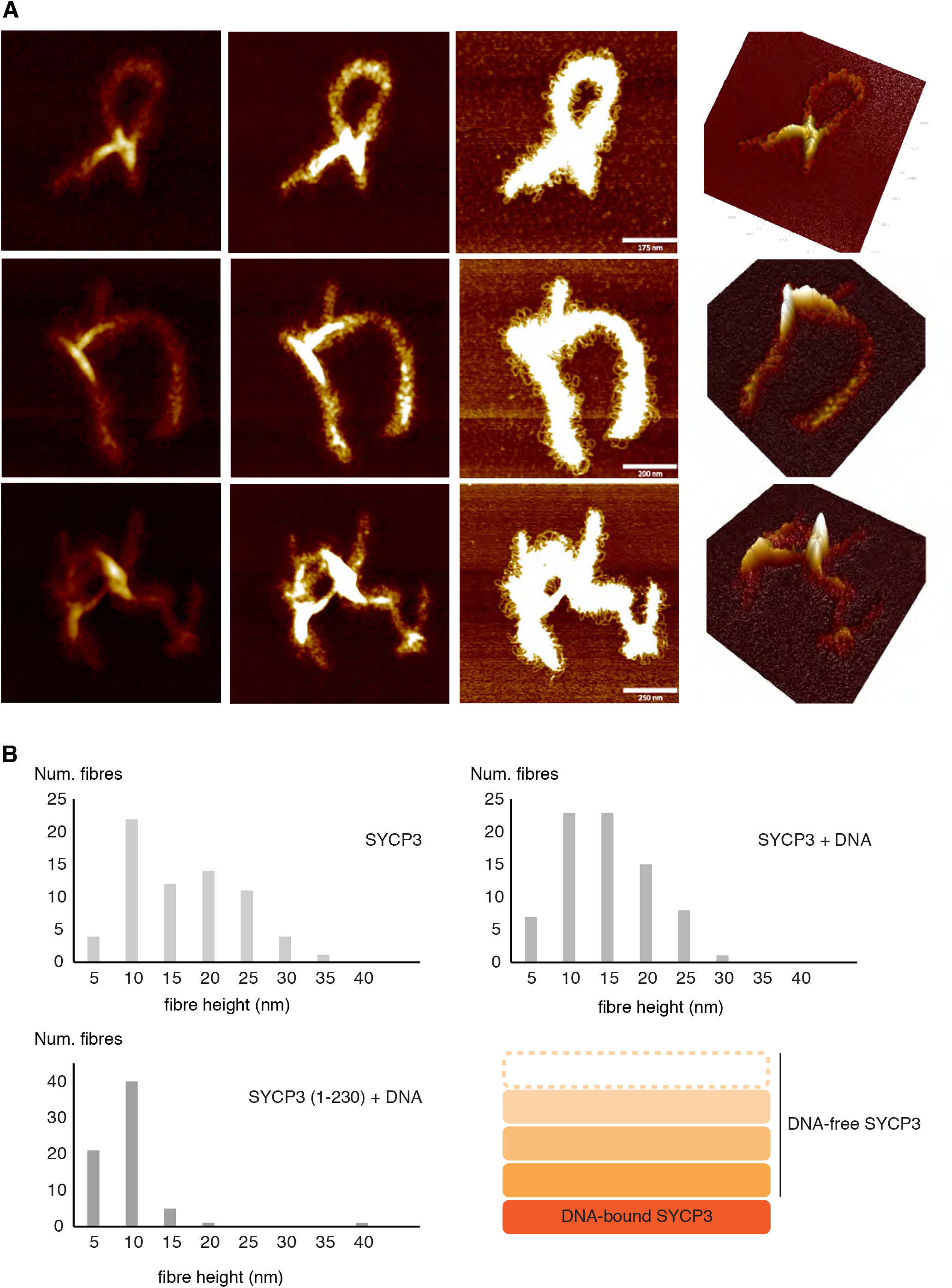
Height distribution of the SYCP3 fibres. **A** Three examples of DNA-bound SYCP3 fibre, displayed at decreasing height threshold from left to right to highlight the variable height of the fibre along its axis. For each example, the right-most panel shows a 3D-representation of the AFM image. **B** Distribution of height values for fibres of DNA-bound SYCP3, DNA-bound SYCP3 1-230 and DNA-free SYCP3. The schematic drawing shows a model for the possible stacking of DNA-free fibres on top of a DNA-bound fibre.

## DISCUSSION

In this paper, we have provided an experimental description of the SYCP3 fibre, in isolation and bound to DNA. We have shown that the filamentous SYCP3 fibres are based on a surprisingly loose and heterogeneous mode of packing, and that the fibres can readily incorporate DNA into their structure. We propose that the combination of self-association and DNA binding, which we have experimentally demonstrated here, constitute the mechanistic basis for the role of SYCP3 in lateral element formation on the chromosome axis.

The homotypic interactions of SYCP3 tetramers within the fibre represent a remarkable example of how a large three-dimensional scaffold can be built rapidly and efficiently out of a small number of homotypic interactions between intrinsically disordered protein regions. Thus, the only apparent inter-particle contacts in the fibre are those between N- and C-terminal tails of the SYCP3 tetramers, leaving the coiled coil core of the SYCP3 tetramer to act as a strut between layers in the fibre. The strut itself appears to be rather flexible, as demonstrated by our EM analysis and the observation of high B factors in the central region of the crystal structure of SYCP3’s helical core (26). The combination of flexible linkages between adjacent SYCP3 molecules and pliability of the SYCP3 helical core endows the fibre with a high degree of plasticity. Thus, rather than a rigid 3D scaffold, the SYCP3 fibre is akin to a flexible chain mail. We note that the peculiar structural properties of the SYCP3 fibre described here seem compatible with the proposed liquid-crystal properties of the SC (32).

The well-documented ability of SYCP3 to form polymeric, fibrous structures, upon heterologous expression in mammalian cells and as a recombinant protein, indicates that the fibrous state represents its functionally relevant form and provides a mechanism for its widespread coating of the chromosome axis. Our AFM experiments provide the first direct evidence for the ability of the SYCP3 fibre to bind DNA. The array of DNA loops protruding from the DNA-bound SYCP3 fibre indicates that, rather than a sporadic phenomenon, the interaction with DNA is a specific property of the SYCP3 fibre. Thus, our findings suggest that spreading of polymeric SYCP3 on the chromosome axis is coupled to its association with chromosomal DNA.

The plasticity of the SYCP3 fibre highlighted by our data suggests that, rather than fulfilling a structural role in determining chromosome architecture, SYCP3 forms a protein layer coating the surface of the chromosome axis (**Fig. 6**). Thus, SYCP3 deposition would take place in the context of the existing chromosome structure, determined by meiotic cohesins. The DNA-bound layer of SYCP3 molecules would then spread over the chromosome surface, achieving a uniform coating at pachytene. The function of such SYCP3 coating might include maintenance of local chromosomal architecture, modulating the availability of defined chromosomal domains during recombination, and acting as a recruiting platform for meiotic factors such as the HORMAD proteins and LE components such as SYCP2. At the end of the meiotic prophase, the SC is rapidly disassembled and SYCP3 disappears from the chromosome axis, except for the centromeric region (33, 34). As formation of the SYCP3 fibre is a fast and irreversible process *in vitro*, SYCP3 removal is likely to require post-translational modifications that weaken its self-association and target it for degradation (35).

**Figure 6.**
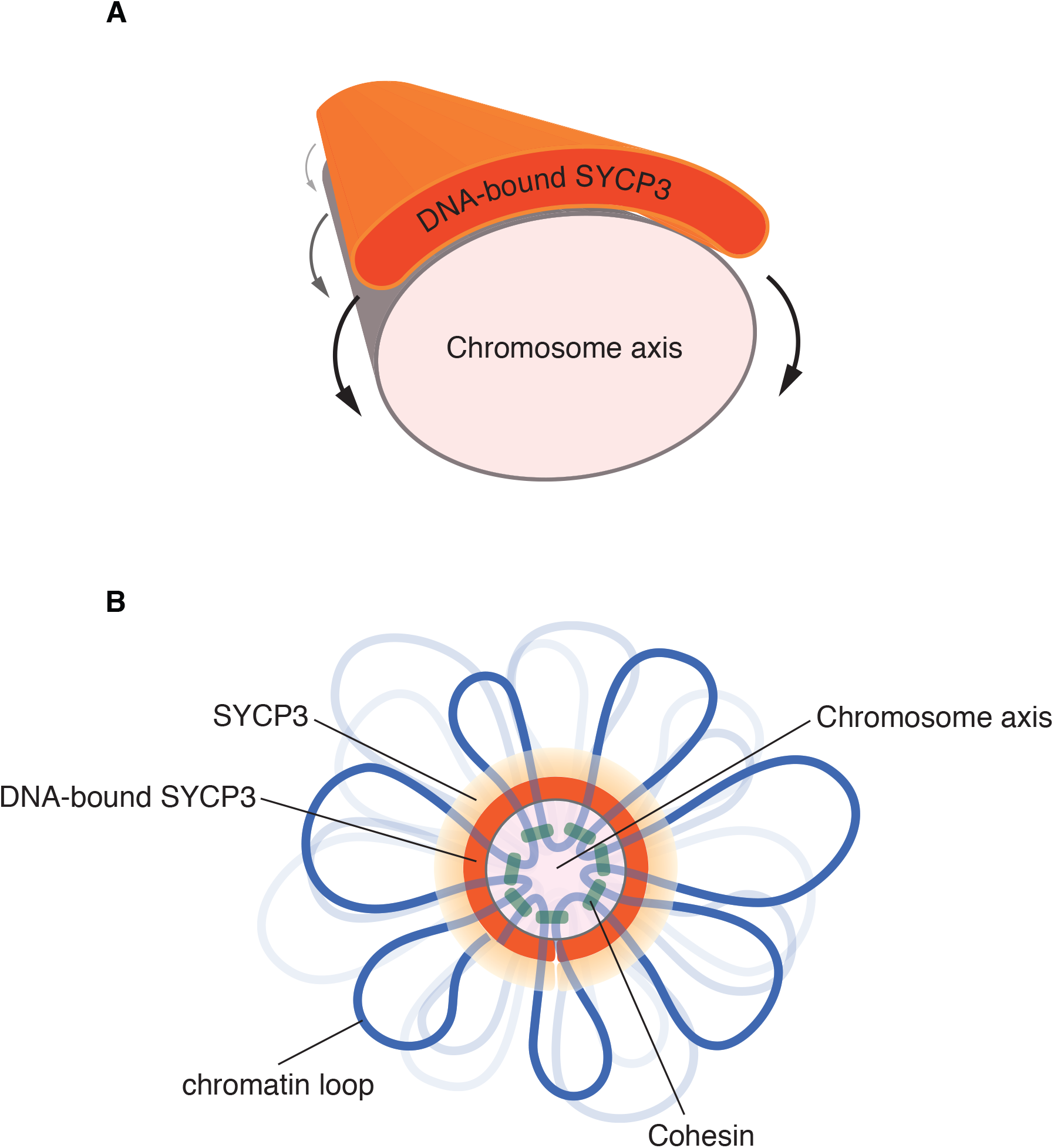
A mechanism of SYCP3 deposition on the meiotic chromosome. **A** Upon accumulation on the chromosome surface, SYCP3 polymerises into a flexible DNA-bound layer that spreads and eventually coats the entire length of the chromosome. **B** Schematic model of the cross-section of the meiotic chromosome axis. SYCP3 coating takes place in the context of the cohesin-dependent structure of the meiotic chromosome, and it is likely to represent the basal level of the lateral element of the synaptonemal complex.

What is the functional relationship between the DNA-free and DNA-bound states of the SYCP3 fibre? A SYCP3 truncation that abolishes fibre formation limits the size of the DNA-bound fibres (**Fig. 4A**), indicating that the DNA-free and DNA-bound states of the fibre share at least some of their homotypic interactions. Conversely, the AFM analysis shows that the DNA-bound fibre lacks the characteristic periodic striations, suggesting that at least some of the underlying interactions between SYCP3 molecules have changed. We did not observe hybrid fibres containing both DNA-bound and DNA-free segments, so their seamless integration might be impossible within the plane of the fibre. However, it is conceivable that DNA-bound and DNA-free forms of the SYCP3 fibre might stack together in a multi-layer structure. Supporting evidence comes from height measurements of DNA-bound and DNA-free fibres (**Fig. 5B**): whereas the height of DNA-bound fibres formed by the truncated SYCP3 defective in self-association is centred around 10 nm, fibres of full-length SYCP3, in both DNA-free and DNA-bound states, show a broader distribution around larger height values. We speculate that the presence of additional SYCP3 on top of a DNA-bound layer might ensure lateral element stability or reinforce the functional insulation of specific chromosomal domains.

Our findings provide a structural and mechanistic basis to help unravel the role of the SC lateral element in meiosis. Future investigations will aim to elucidate the interaction mechanism of the SYCP3 fibre with DNA and with other known components of the meiotic chromosome axis. Such studies will be essential to improve our knowledge of the molecular mechanisms underlying meiotic recombination and the impact on fertility when such mechanisms become defective. The realisation that meiotic gene products, including SYCP3, are upregulated in a number of cancers (36, 37) further highlights the importance of these studies to understand the pathological mechanisms that contribute to genomic instability in human cells.

## METHODS

### Protein expression

Amino acid sequence corresponding to 1-236 (full length) and 1-230 of human SYCP3 were cloned into the pHAT4 vector (38) for expression in bacteria with an N-terminal TEV-protease cleavable His_6_-tag. Recombinant proteins were expressed in Rosetta 2 (DE3) *E. coli* (Novagen). Transformed cells were plated out on LB agar supplemented with 34 μg/ml chloramphenicol and 100 μg/ml ampicillin. Bacterial colonies were transferred to 1 litre 2xYT (100 μg/ml ampicillin, 34 μg/ml chloramphenicol) at 37°C and grown until 0.6 OD600 in a baffled two litre flask. Bacteria were then induced (0.5 mM IPTG, 25°C) and grown overnight, harvested by centrifugation (4000*g*, 20°C, 20 min), resuspended in 20 mM Tris-HCl pH 8.0, 400 mM KCl buffer (25ml buffer per pellet of 1 litre culture) containing EDTA-free protease inhibitors (Sigma) and stored at −80°C.

### Protein purification

The bacteria were lysed by sonication, the lysate was clarified by centrifugation (30000*g*, 4°C, 30 min) and passed through a 45 μm filter. SYCP3 proteins were first purified using Ni-NTA agarose resin (Qiagen). A Ni-NTA column (2 ml) was equilibrated with five column volumes of wash buffer (20 mM Tris pH 8.0, 400 mM KCl). The clarified lysate was passed over the column under gravity flow. The column was washed with 20 column volumes of wash buffer to remove unbound bacterial proteins, followed by 4 columns volumes of wash buffer with 20 mM imidazole to remove weakly binding bacterial contaminant proteins. Bound SYCP3 was recovered in two elution steps, using wash buffer with 100 mM and 200mM imidazole. The 200 mM imidazole elution typically contained the cleanest recombinant protein, and was purified further as described below.

#### Full-length SYCP3

To cleave the His_6_-tag, TEV protease was added at mass ratio of 1:50 and incubated at 4°C overnight. To help remove nucleic acid contamination present after the Ni-NTA step, benzonase was also added during the TEV treatment. Full-length SYCP3 was then concentrated by centrifugal device and further purified by gel filtration over a Superdex 200 16/60 column equilibrated in 20 mM Tris-HCl pH 8.0, 400 mM KCl. Peak fractions were analysed by SDS-PAGE, pooled, spin concentrated (4500*g*, 10°C) using 4 ml Amicon Ultra 10,000 MWCO filter units (Millipore), flash frozen and stored at −80°C.

#### SYCP3 1-230

To cleave the His_6_-tag, TEV protease was added at mass ratio of 1:50 and incubated overnight on ice. SYCP3 1-230 was further purified by cation-exchange over a 5-ml HiTrap SP column (GE Healthcare) and Heparin Sepharose chromatography (GE Healthcare), using a linear salt gradient from 150 mM to 1M KCl gradient in 20 mM Tris-HCl pH8.0, 150 mM KCl. Peak fractions were pooled, spin concentrated (4500*g*, 10°C) using 4 ml Amicon Ultra 10,000 MWCO filter units (Millipore), flash frozen and stored at −80°C.

### Salt-dependent aggregation of SYCP3

The aggregation index (AI) of the SYCP3 protein was analysed adapting a published protocol(39), using the following formula:

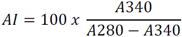

Where A340 and A280 are the absorbance of the SYCP3 sample at 340 nm and 280 nm, respectively. The AI provides an indication of the aggregation state of a protein sample in solution, independent of its concentration. The AI was used to study the salt dependency of SYCP3 aggregation, measuring five SYCP3 samples at 25 μM in 20mM TrisCl buffer pH 8.0 and KCl concentrations ranging from 100 to 500 mM.

### Sample vitrification for cryo-electron tomography

100 μl of BSA-coated 10 nm gold bead suspension (Aurion, Wageningen, The Netherlands) were centrifuged at 14000*g* for 10 min, the clear supernatant discarded, and the gold beads resuspended in 50 μl ice-cold 20 mM Tris, pH 8.0 (DB buffer). Quantifoil grids (Cu 200 mesh, 2/1 holey carbon, Quantifoil Micro Tools GmbH, Jena, Germany) were plasma-cleaned for 30 s. Vitrification was carried out in a Vitrobot Mk IV (FEI / Thermo Fisher, Eindhoven, The Netherlands), with the sample chamber set to 4 °C and 90% humidity, a blotting strength parameter set to “10” and a blotting time of 5 s. 3 μl of previously prepared gold particle resuspension were applied to the grid. The droplet was allowed to dry for about 2 min, until only a thin liquid film covered the grid. Immediately afterwards, 4 μl of ice-cooled SYCP3 protein solution (14 mg/ml), freshly mixed 1:1 with DB buffer, to reduce salt concentration and promote fibre formation, were pipetted onto the grid. After a short incubation time of 10 s, the grid was transferred to the Vitrobot, blotted and plunged into a liquid nitrogen-cooled bath of 37 % ethane, 63 % propane.

### Cryo-electron tomography of SYCP3 fibres

Ten tilt series of fibres were recorded with a Tecnai G2 Polara (Thermo Fisher), equipped with a 300 kV FEG, a Gatan post-column energy filter and a K2 Summit direct electron detector (Gatan). Images were recorded at a working magnification factor of 34,000×, a pixel size of 3.45 Å/px and a dose rate of 0.8 e/Å^2^/s with a total dose per tilt series of 70 – 100 e/Å^2^. The target defocus was 2.5 μm. For each tilt series, 61 micrographs were recorded at 2° increments with a tilt order of −30° to +60° followed by −32° to −60°.

### Tomographic reconstruction and subtomogram analysis

Here we provide a complete description of the subtomogram analysis of the SYCP3 fibres under native conditions. Further information on the method can be found in our recently published protocol for analysis of filamentous protein assemblies such as the SYCP3 fibres (40). Drift correction for the tilt series micrographs was performed with MotionCor2 (41). The tomograms were backprojected in IMOD (42), using the embedded 10 nm gold markers as fiducials for the alignment. Due to the characteristically low signal-to-noise of the tomographic reconstructions, it was not practical to identify and select individual SYCP3 tetramers as subvolumes. Instead, subvolume coordinates were picked along the striation density of the fibres. To facilitate the detection of the striations within the fibre, the tomographic volumes were low-pass filtered with a cut-off at 0.44 nm^-1^. Two coordinate models were created using IMOD (42), by manually tracing the ends of individual striations with a single contour and then interpolating equidistant points within each contour via the program addModPts of the PEET package (43, 44). The first model featured a smaller set of 5756 coordinates, widely spaced at 44 nm (128 voxels), which was used to generate an initial reference structure *de novo* (see below). The second model comprised a total of 46051 subvolume coordinates, at a narrow spacing of 16.5 nm (48 voxels) in the XY plane and 22 nm (64 voxels) along Z. This set of coordinates was used later for masked structural refinement.

Subtomogram averaging was performed in Relion 2.1, adapting a modified protocol by Bharat and Scheres (45). The IMOD coordinate models were converted to ASCII format using model2point (42). CTF estimation was performed with ctffind4 (4.0.15) (46), via the relion_prepare_subtomograms.py script(45). The first set of subvolume coordinates was used to extract 5767 non-overlapping subvolumes, rescaled from 128^3^ voxels unbinned to a size of 64^3^ voxels (equivalent to 44×44×44 nm^3^ at a pixel size of 6.9 Å/pixel). An identical set was extracted and resized with a binning factor of 2. Due to the manual picking of coordinates, each subvolume was only roughly centred on a striation density. Since the long axis of the fibres was oriented parallel to the grid surface, a translational alignment in the XY plane was sufficient to precisely align all subvolumes within the fibre pattern. For this, Z-projections of all rescaled subvolumes were used as the input for a reference-free 2D classification of three classes. This step served as an XY-plane alignment measure, as well as a screening step to remove unsuitable particles, such as subvolumes picked too close to the edge of a fibre, from the following processing steps. The majority of particles (51%) contributed to a class that showed a clear striation density layer at its centre, with separate elongated densities of around 2×20 nm in size connecting orthogonally to adjacent striations, in good agreement with the dimensions of the coiled-coil core region of an SYCP3 tetramer (26). Using the relion_2Dto3D_star.py script (45), the file paths of 2D projections contributing to this class average were translated back to their respective 3D subvolumes in a new particles_subtomo_2Dto3D.star table. In addition to the file paths, the translational shift columns (_*rlnOriginX* and _*rlnOriginY*) were added for each subvolume. Using this list of XY-centred subvolumes, a reference-free 3D angular search was performed. The resulting single-class average constituted a low resolution *de novo* structure and was rotated with IMOD’s rotatevol function (http://bio3d.colorado.edu/imod/doc/man/rotatevol.html) so that the striation plane was orthogonal to the Y axis of the subvolume. The rotated structure was subsequently refined in 3D classification and 3D auto-refinement steps against the entire first set of subvolumes. The refined structural average featured a clearly aligned centred striation density, with orthogonally connecting elongated SYCP3 tetramers, roughly arranged in a rectangular mesh along the striation plane.

Using the second set of coordinates, 46051 overlapping subvolumes of 44×44×44 nm^3^ (128^3^ voxels at the unbinned pixel size of 3.45 Å/px) were extracted and rescaled by a factor of 0.5, corresponding to a subvolume size of 64^3^ voxels at twice the pixel size (6.9 Å/px). This larger set of subvolumes was used for a masked 3D classification, with the refined *de novo* structure created in the previous step serving as the input reference. The reference mask was created in Matlab/Tom Toolbox (47), and consisted of a narrow cylinder of 24 voxel diameter (equivalent to 16.5 nm at a binning factor of 2) with its long axis oriented parallel to the Y axis and a smooth gaussian falloff within a rescaled 64^3^ voxel volume. The masked 3D classification resulted in 4 class averages, with nearly equal class distributions. Subvolumes with a large translational shift (with individual translational shifts in X, Y and 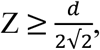 *d* being the distance of subvolume coordinates in voxels) were removed to exclude auto-correlation effects of potentially overlapping subvolumes. Each trimmed class data set was then 3D auto-refined.

These final refined class averages showed a high degree of heterogeneity in the organisation of the SYCP3 tetramer within the fibre, with resolutions of around 30 - 27 Å. Additional hierarchical 3D classification of these classes, focusing on a single striation-striation repeat by using a reference mask of only a half cylinder and splitting each refined subvolume subset into 8 classes, resulted in a clearer separation of structural motifs but did not improve resolution. Consistent with this, the class distribution was close to 12.5% (±2 %) for the refined subsets 1, 2 and 3, confirming the presence of high structural heterogeneity; subset 4 showed a slight deviation from this pattern as it featured two heavily warped class averages with 16.6% and 15.9% subvolume distribution, probably due to the missing wedge-induced elongation of subvolumes, which could not be compensated for with sufficient precision.

### Atomic force microscopy

For analysis of the DNA-free fibres, purified SYCP3 was diluted to micromolar concentrations into a low-salt buffer (20mM TrisCl pH 8.0, 150 mM NaCl) to induce fibre formation, deposited on freshly cleaved mica and incubated at room temperature for 10 minutes. For fibre formation in the presence of DNA, 0.5 nM plasmid DNA was mixed with a 5-fold molar excess of SYCP3 in the low-salt buffer shortly before deposition on mica. The majority of experiments were performed with pUC19 DNA, but other circular double-stranded DNA such as PhiX174 RFII were tested with similar results. The samples were washed five times with BPC water (Sigma) and dried under a gentle nitrogen stream. Imaging was performed using a Bruker Dimension FastScan AFM, in air, at ambient temperature. The probes used were FastScan-A (Bruker), with resonant frequency of 1400 kHz and the drive frequencies were approximately 5% below the maximal resonance peak. AFM images were acquired at a rate of four frames per minute and plane-fitted to remove tilt. Each scan line was fitted to a first-order equation, using the NanoScope analysis software (Bruker).

## ACKNOWLEDGMENTS

We would like to thank John Heumann and Johanna Syrjanen for their comments and advice. We would also like to thank Joseph Maman for advice and assistance with the SYCP3 fibre formation assay. This work was supported by a project grant from the Medical Research Council (MR/N000161/1) to LP.

## AUTHOR CONTRIBUTIONS

DB and LR expressed and purified SYCP3. D.B. collected the cryo-electron tomography data, assisted by JMP, and performed the sub-tomogram averaging analysis of the SYCP3 fibre. LR and IM prepared the samples for atomic force microscopy and IM performed the imaging and analysis of the data, with help and advice from RMH. DB, LR, IM and LP designed experiments and analysed results. LP conceived the project and directed the research.

## Supplementary Figure legend

**Supplementary figure 1**. Aggregation Index (AI) analysis of the salt-dependency of SYCP3 fibre formation. The AI is measured spectrophotometrically using the formula in the figure, in the presence of decreasing concentration of KCl, to define the concentration of salt leading to fibre formation.

**Supplementary figure 2**. Representative electron micrographs of native SYCP3 fibres in vitreous ice, prepared as described and shown in Figure 1.

**Supplementary figure 3**. Generation of coordinates models for subtomogram extraction. **A, B** Each tomogram was sectioned along the Z-axis to generate slices in the XY plane at a distance of 64 pixels (22 nm) in Z. **C** A low-pass filter was applied to boost the signal of the striation pattern. **D** At every 64^th^ XY-slice along the fibres with discernible striations, a two-point contour was placed along each striation. **E** Additional points were added at a constant interval to the contours as detailed in the Methods. Two coordinate models were generated per reconstructed tomogram: the first model featured a wide spacing of 44 nm (128 voxels) between subtomograms; the second model had a narrower coordinate spacing of 16.5 nm (48 voxels.

**Supplementary figure 4**. Workflow for the process of sub-tomogram averaging.

**Supplementary figure 5**. Further hierarchical classification of the final 4 class averages, shown in Figure 2, did not improve resolution or lead to the identification of distinct structural conformations within the fibre.

**Supplementary figure 6**. Two examples of AFM images of SYCP3-DNA fibres. The panels on the right show an enlarged section of the image, to highlight the features of the DNA-bound protein structure.

**Supplementary figure 7.** AFM images of circular and linearized pUC19 DNA.

**Supplementary movie 1**. Reconstructed cryo-electron tomograph of a SYCP3 fibre in vitreous ice.

